# Chronic evoked seizures differentially worsen cognitive function and long-term neuropsychiatric comorbidities in Alzheimer’s disease-related genotypes

**DOI:** 10.64898/2026.11.20.768142

**Authors:** Kevin M. Knox, Stephanie Davidson, Jennie Fang, Melissa Barker-Haliski

**Author notes:** corresponding author, Address for correspondence: 1959 NE Pacific St. Box 357610, Seattle WA 98195 USA.

## Abstract

Variants in one of three deterministic risk genes - amyloid precursor protein (APP), presenilin 1 (PSEN1), or presenilin 2 (PSEN2) – lead to early-onset Alzheimer’s disease (EOAD), which also carries a high risk of spontaneous seizures. Seizures may worsen AD-related behavioral comorbidities, including depression, anxiety, and cognitive impairment. Using the corneal kindling model of secondarily generalized focal seizures in two distinct aged male and female EOAD-associated mouse models, we assessed the interaction between investigator-initiated chronic seizures and EOAD genotype on AD-related behavioral comorbidities. Genotypes were selected to model amyloid-β independent and dependent AD risk factors and included: PSEN2-N141I (amyloid-β independent model) versus wild-type (PSEN2 control) mice, and APP/PS1 (amyloid-β dependent model) versus wild-type littermates (Tg-). Sham and fully kindled mice then underwent behavioral assessment in the saccharin preference test of anhedonia-like behavior, the Barnes maze test of spatial learning and memory, and a 67-hour home cage monitoring of circadian activity. EOAD genotype only significantly affected kindling acquisition of aged APP/PS1 females. However, there was no significant genotype or kindling effect on saccharin preference, suggesting no adverse impact on anhedonia-like behavior. Cognitive performance in the Barnes maze was significantly worsened by seizures exclusively in APP/PS1 female mice. However, seizures generally worsened Barnes maze performance of male PSEN2-N141I and PSEN2 control mice. Finally, chronic seizures consistently worsened exploratory behavior patterns during home cage circadian activity monitoring exclusively in APP/PS1 male and female mice. This study reveals the AD-related heterogeneity of seizure-evoked behavioral consequences on spatial working memory and circadian behaviors.

## Introduction

Patients with Alzheimer’s disease (AD) have no medical options that can safely slow disease progression. Current first-line therapeutics for AD are cholinesterase inhibitors and the NMDA antagonist memantine (with or without a cholinesterase inhibitor), neither of which alter the underlying disease processes or long-term cognitive burden. More recently approved monoclonal antibodies that target amyloid-β (Aβ) carry significant adverse events liability, thus leaving AD patients and their caregivers in need of novel disease-modifying interventions. AD is increasingly recognized to be associated with undetected (and thus untreated) comorbid seizures ^1^, which potentially hasten functional decline and exacerbate AD-related behavioral comorbidities ^2; 3^. Silent hippocampal (i.e., focal) seizures are documented in AD^1^, implicating excessive network hyperexcitability as an underexplored contributor to the behavioral sequelae of AD. For example, neuropsychiatric comorbidities of AD (depression, anxiety, sleep disruption^4^) are detrimental to AD patient quality of life and people with epilepsy also often experience these same comorbidities that also worsen quality of life^5; 6^. Sleep-wake disturbances in both AD and epilepsy may exacerbate cognitive deficits and worsen neuropsychiatric disturbances. For example, a small study of patients with advanced AD reported that those with new onset seizures had faster declines in language assessment.^7^ Seizures in dementia patients can significantly shorten lifespan versus that of unaffected individuals^8^. Unmitigated neuronal hyperexcitability is thus an underexplored mechanism that likely worsens the functional and pathological burden of AD.

Most mechanistic evidence on the presentation of and impact from seizures in AD has come from clinical^1; 9–11^ and preclinical work with amyloid-precursor protein (APP)-overexpressing mouse models ^12–16^. However, mutations in presenilin (PSEN) 1 and 2 also lead to rare, but Mendelian, forms of early-onset AD (EOAD). Notably PSEN2 variants may even be masked in the late-onset AD population^17; 18^ or potentially misdiagnosed as frontotemporal dementia ^19^, providing an important and differentiated opportunity to assess the degree to which seizures impact disease progression and outcomes across the spectrum of AD and AD-related dementias. The most common AD-related PSEN2 variant, the Volga German N141I^20^, leads to clinical seizures ^20–22^. However, PSEN2 variant mouse models do not routinely demonstrate amyloid-β (Aβ) plaques albeit they do show age-related pathology and aberrant neuroinflammation and microgliosis^23–25^. Seizures in human epilepsy also heighten neuroinflammation and microgliosis ^26; 27^, thus our present study hypothesized that PSEN2-N141I variants may increase seizure susceptibility through an amyloidogenesis-independent mechanism.

Most prior seizure studies in AD-related mouse models have been largely limited to using long-duration monitoring of infrequent spontaneous recurrent seizures (SRS) in single-housed mice. Yet, social isolation stress can adversely affect seizure susceptibility in rodents ^28^, which may negatively increase disease burden in already seizure-prone AD models. Unlike the long-duration recording of SRS in single-housed mice,^12; 15; 16; 29^ corneal kindling is a routinely used rodent acquired temporal lobe epilepsy (TLE) model ^30; 31^ that supports long-duration social housing ^32; 33^ and reproduces the key clinical TLE features, including neuroinflammation^34^ and behavioral deficits ^35; 36^. Our present study thus directly assessed how experimenter-controlled chronic seizure history with the PSEN2-N141I variant vs APP overexpression (APP/PS1 mice) could impact behavioral comorbidities of AD in late life. Our primary study objective was to directly demonstrate whether chronic secondarily generalized focal seizures incite and/or aggravate neuropsychiatric symptoms, cognitive deficits, and neuropathology, rather than merely demonstrating that infrequent SRS occur rodent AD models that also exhibit behavioral and cognitive deficits in late life.

Pharmacological suppression of focal seizures in patients with AD using antiseizure medicines (ASMs) represents a novel disease-modifying strategy to slow AD progression and potentially reduce the severity of neuropsychiatric comorbidities ^11^. Seizures may be a manageable feature of AD ^37–39^ because seizures in older adults are generally not treatment-resistant ^40^ and numerous ASMs are already FDA approved and frequently used in older adults with epilepsy^41^. Yet whether AD-related risk factors differentially interact with seizures to impact disease trajectory is less clear. Both APP/PS1 and PSEN2-N141I variant mice show unique and distinct pharmacological response to the most commonly prescribed ASMs for older adults with epilepsy ^24^, as well as exhibit differential outcomes following evoked focal seizures at presymptomatic ages ^42^. Accordingly, AD-related risk genes may also differentially interact with chronic seizures to modify discrete behavioral domains that influence neuropsychiatric and cognitive burden in late-life. It is also unclear whether seizures in AD, in fact, directly aggravate the behavioral burden to any differential degree in male versus female AD models. As a result, a second major goal of this present study was to establish the direct contribution of chronic seizures to the burden of AD-related neuropsychiatric comorbidities, cognitive function, and neuropathology in aged rodents so that future preclinical and clinical interventions can be more rationally selected for seizures in older adults with AD.

## Materials and Methods

### Animals

For the initial studies to establish the age-related changes in neuronal hyperexcitability and focal seizure susceptibility across the lifespan (Figure 1 and Figure 2), male and female PSEN2-N141I and APP/PS1 mice were tested between age 2-12 months (Figure 1). For the corneal kindling studies (Figures 3-7), male and female PSEN2-N141I and APP/PS1 mice were then tested between age 10-20 months. For all studies, mice were housed on a 14:10 light cycle (on at 6h00; off at 20h00) in ventilated cages with free access to food and water as previously described ^43^. All behavioral testing was performed between the hours of 7h00 and 19h00. All animal work was approved by the University of Washington Institutional Animal Care and Use Committee (protocol 4387-01) and conformed to ARRIVE guidelines ^44^. Behavioral seizure induction and comorbidities testing in all studies was performed blinded to genotype. The PSEN2-N141I and PSEN2 control mouse lines were derived and maintained as previously described ^21; 24^. The APP/PS1 mouse line was maintained as previously described^24^, with non-transgenic littermates serving as the APP control mice.

**Figure 1.**
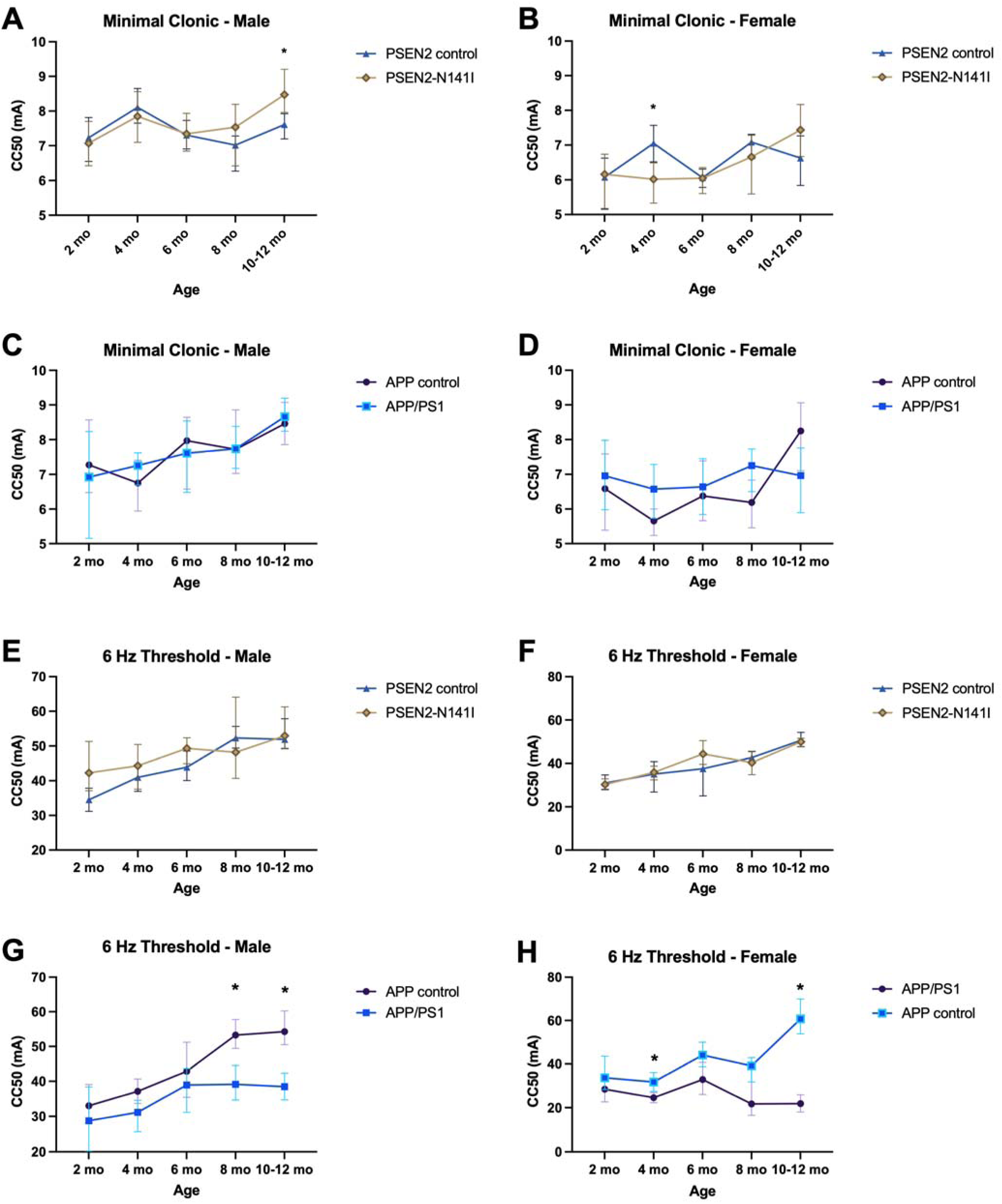
Threshold to an evoked seizure using either a 60 Hz 1-10 mA 0.2 second stimulation (minimal clonic) or 6 Hz 20-80 mA 3 second stimulation (6 Hz) generally increases with advanced age in male and female micewith and without AD-related risk genes, which is blunted around the time that Aβ deposition begins in APP/PS1 mice (typically 4.5-6 months-old). A-D) The minimal clonic test is an electrically evoked focal seizure model that is characterized by rhythmic face and forelimb clonus, ventral neck flexion, and may include rearing and falling. A-B) In male and female PSEN2-N141I and matched control mice, there are modest age-related differences in minimal clonic seizure threshold across the lifespan up to 12-months-old. C-D) In male and female APP/PS1 and matched control mice, there were no significant differences in minimal clonic seizure threshold up to 12-months-old. E-H) The 6 Hz seizure threshold test is an electrically evoked focal seizure model that is characterized by vibrissae twitching and forelimb clonus, which activates distinctly different brain regions than the minimal clonic test. E-F) In male and female PSEN2-N141I and matched control mice, there were no notable differences in age-related seizure threshold changes up to 12-months-old. G-H) However, in male and female APP/PS1 and matched control mice, there were significant differences in evoked seizure threshold as early as 8-months-old in males (G) and as early as 4-months-old in females (H). These findings suggest that APP/PS1 mice lose the normal age-related increase in seizure threshold, likely coinciding with the onset of Aβ plaque deposition. Data are expressed as median convulsant current (CC50) with 95% confidence intervals, as determined by a probit regression model of Finney. * indicates non-overlapping 95% confidence intervals, reflective of statistically significant differences.

**Figure 2.**
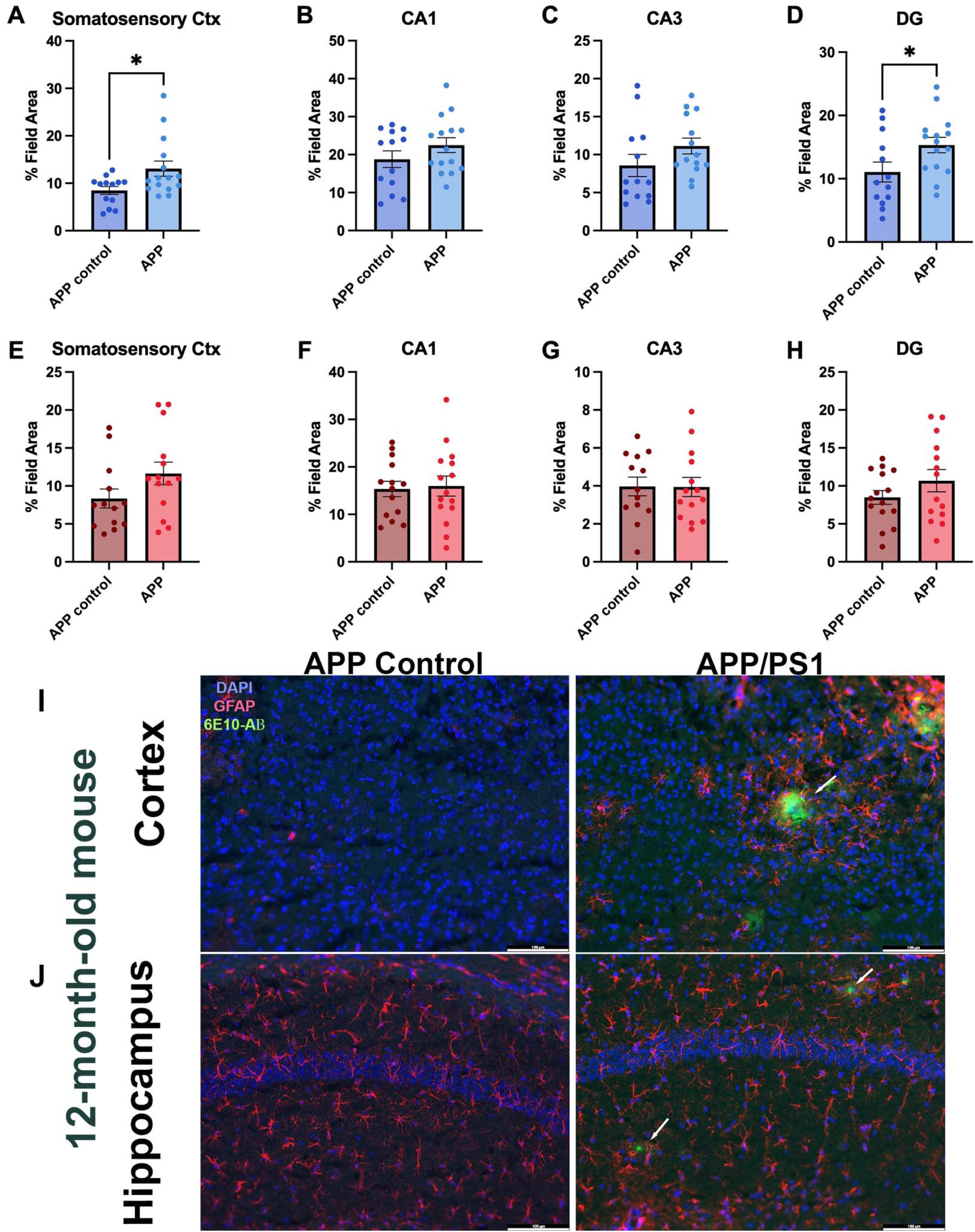
Aged (12-month-old+) APP/PS1 mice from the seizure threshold studies show increased EAAT2, but not GAD65/67, expression and Aβ plaque deposition in cortex and hippocampus at 12-months-old. A-D) Male and female APP/PS1 mice were euthanized after the completion of minimal clonic and 6 Hz seizure threshold testing for ex vivo analysis of EAAT2 expression across several brain regions involved in neuronal hyperexcitability. There were significant increases in EAAT2 expression in APP/PS1 mice in A) somatosensory cortex (t=2.548, p=0.019) and D) dentate gyrus (DG; t=2.145, p=0.043) of hippocampus, but not in other hippocampal subregions. E-H) However, there is no change in GAD65/67 expression in these same brain regions, suggesting no global shift in inhibitory neurotransmission. I) Further, Aβ plaques in cortex and hippocampus of 12-month-old APP/PS1 mice were surrounded by regions of reactive astrocytes (as demonstrated by GFAP immunoreactivity), but plaques were not evident in the brains of control mice, confirming development of AD-related neuropathology in this mouse model. These seizure threshold data and AD-related neuropathological hallmarks compelled further study of chronic seizure-related functional and behavioral effects in late life in two AD mouse models. * indicates p<0.05, as measured by t-test.

**Figure 3.**
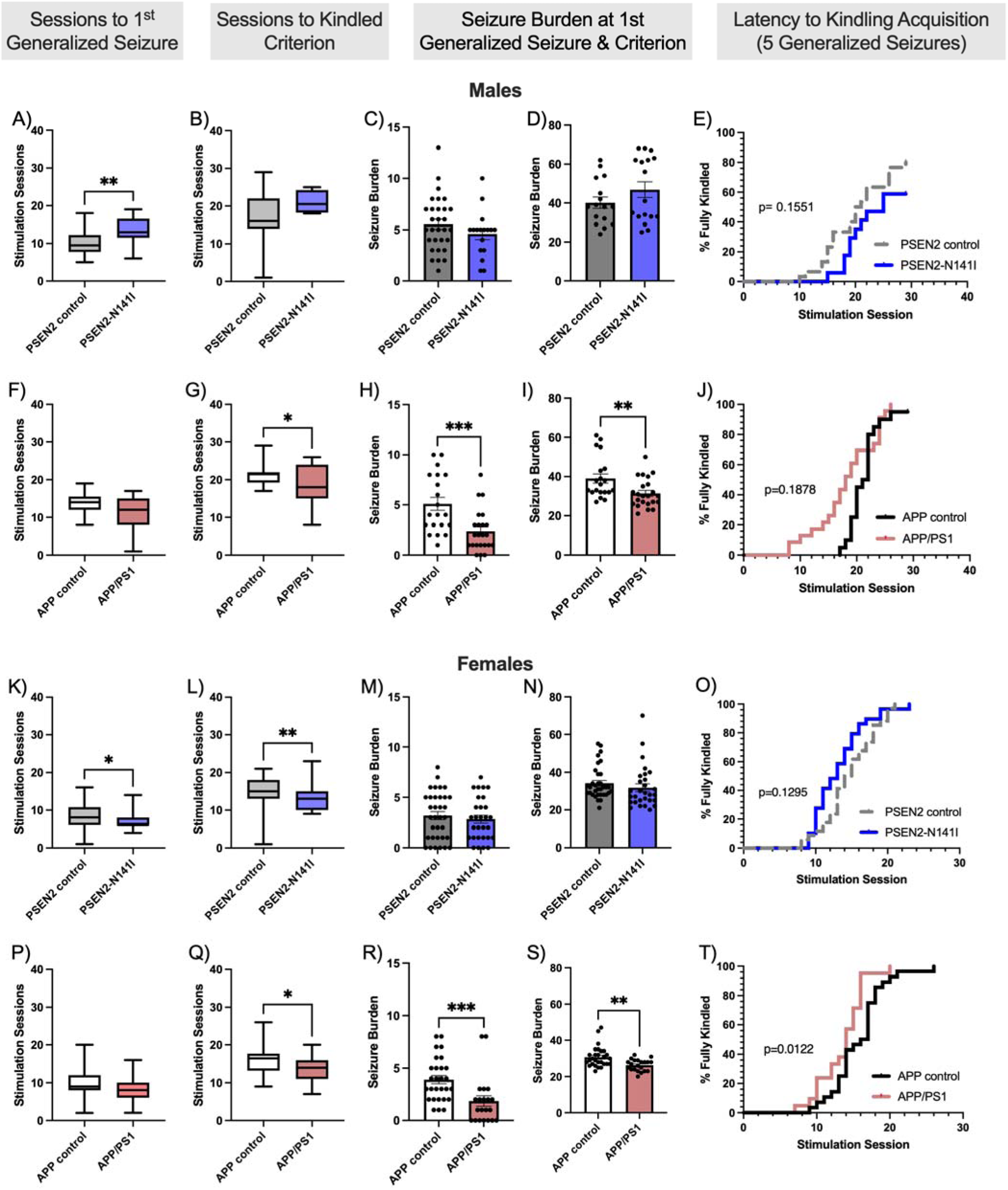
Aged female, but not male, mice with AD-related genetic risk factors showed increased susceptibility to developing a hyperexcitable neuronal network, as measured by propensity to acquire the fully kindled state, modeling acquired epilepsy. Following twice-daily corneal stimulation with a 3 sec 1.6-3.0 mA 60 Hz electrical current, male and female PSEN2-N141 and APP/PS1 mice and their matched controls were tracked for the number of sessions to experience a first secondarily generalized seizure (A, F, K, and P). PSEN2-N141I transgenic male mice showed significant delay to experience a first convulsive seizure (A; Mann-Whitney U = 131.5, p=0.0053), whereas PSEN2-N141I females showed a modestly faster latency to experience the first seizure (K; Mann-Whitney U = 466.5, p=0.015). The number of sessions required for animals to attain fully kindled criterion, as defined as five consecutive Racine stage 5 seizures, was similarly tracked across all sexes and genotypes (B, G, L, and Q) with APP/PS1 male and APP/PS1 female mice consistently showing faster kindling acquisition (G, Mann-Whitney U = 148, p=0.045; and Q, Mann-Whitney U = 186, p=0.028), but only PSEN2-N141I females showed faster kindling rates versus matched controls (L, Mann-Whitney U = 447, p=0.008). The seizure burden for all groups was similarly assessed (C-D; H-I; M-N; and R-S), which is a measure of seizure frequency and severity over a monitoring period. Only male and female APP/PS1 mice showed differences in seizure burden at the first generalized seizure presentation and at the end of kindling (H-I; R-S); PSEN2-N141I mice did not differ from their matched controls. Finally, the latency to kindling acquisition, defined as five consecutive Racine stage 5 seizures, was tracked in all mice (E; J; S; T) and analyzed by Kaplan-Meier plot. Only aged female APP/PS1 mice showed a significantly increased latency to kindling acquisition relative to matched controls (X^2^=6.29, p=0.012). * = p<0.05, ** = p<0.01, *** = p<0.001.

### Acute seizure threshold testing

Focal seizures have been documented in clinical AD cases ^1^, thus we first used two distinct rodent focal onset seizure induction models^45^ to define the age-related changes in neuronal network integrity and connectivity in awake, freely-moving PSEN2-N141I and APP/PS1 mice from young adulthood (2 months) to an aged state (12-months-old). Discrete electrical stimulation protocols can differentiate limbic, focal impaired awareness (6 Hz) and forebrain (minimal clonic) seizure thresholds in rodents^45^. Minimal clonic seizures are characterized by rhythmic face and forelimb clonus, ventral neck flexion, and may include rearing and falling. Unlike a maximal electroshock seizure (MES;^46^), the minimal clonic test does not induce tonic hindlimb extension ^47–50^ or seizure-induced death ^51^, making it well-suited to longitudinal seizure threshold monitoring. A 6 Hz seizure is characterized by forelimb clonus, vibrissae twitching, behavioral arrest, and Straub tail ^52; 53^. The minimal clonic and 6 Hz seizure thresholds were determined in male and female AD and control mice aged up to 12-months-old according to the staircase procedure ^54^. Briefly, an electrical current was delivered bilaterally to anesthetized (0.5% tetracaine) corneas of lightly restrained mice (held in-hand) for the specific acute seizure induction protocol (minimal clonic or 6 Hz). A mouse was considered to have had a seizure if it presented with the model-specific behavioral seizure endpoint. The seizure result of each mouse dictated the stimulation current delivered to the next mouse in each group until at least 5 current groups were established between the limits of 0% and 100% of mice with seizure to quantify median convulsant current (CC_50_) and 95% confidence intervals by the probit method of Finney ^55^. Age- and genotype-dependent changes in threshold were determined at 2-, 4-, 6-, 8- and 10-12-months old, with 1-week between tests at each age range. Animals were randomized to stimulation current group during each session for each seizure test. A minimum of n=4 mice/sex per electrical current was tested per seizure threshold test (Figure 1), and group sizes/sex/genotype were: male PSEN2 control = 48; male PSEN2-N141I = 28; female PSEN2 control = 25; male PSEN2-N141I = 40; male APP/PS1 control = 34; male APP/PS1 = 34; female APP/PS1 control = 31; female APP/PS1 = 27.

### Corneal Kindling

Once seizure threshold was established, a separate cohort of mice/genotype were kindled by applying an electrode bilaterally to anesthetized corneas and administering a current (60 Hz, sinusodial pulse; 3.0 mA for male mice, 2.6 mA for female mice) for 3 seconds^24; 42; 56^. Mice were kindled across six separate cohorts, with specific cohorts being subsequently tested in different behavioral testing paradigms.

Mice received maintenance stimulations on days prior to the Barnes maze to sustain chronic seizures, however animals did not receive stimulations on behavioral testing days to separate behavioral performance from the effects of an acute seizure. Sham kindled mice were “kindled” alongside the corneal kindled mice. The sham mice were handled the same way as the kindled mice, except no electrical stimulation was delivered to the cornea during acquisition. Male group sizes were: PSEN2 control = 49 kindled; PSEN2-N141I = 27 kindled; APP control = 22 kindled; APP/PS1 = 26 kindled. Female group sizes were: PSEN2 control = 60; PSEN2-N141I = 38; APP control = 33; APP/PS1 = 25.

### Duration of the Fully Kindled Seizure

After acquiring the fully kindled state, all corneal kindled AD and transgenic control mice were allowed a 3-7-day stimulation-free period. Kindled secondarily generalized seizure duration (Racine stage 4 or greater) was then recorded from all fully kindled mice and compared across genotypes in each sex (supplemental Figure 1), as previously described^56^.

### Open Field Test

Exploratory and anxiety-like behavior was assessed in fully kindled PSEN2-N141I mice only using the open field (OF) task as previously described^36^. Mice were randomized to one of seven OF chambers with their exploration recorded for 10 minutes, as published kindled PSEN2 knockout mice ^57–59^. Mice were placed into the center of the OF chamber and their total distance traveled (cm), vertical activity counts, and horizontal activity counts were measured for 10 minutes and recorded (Accuscan, Omnitech). OF chamber exploration and time spent in the OF center versus the periphery were the primary endpoints.

### Saccharin Preference Test

Anhedonia-like behavior was assessed over 48 hours in sham and fully kindled mice with the saccharin preference test^60^. Mice were single housed for the test duration with *ad libitum* access to bottles filled with either saccharin or water. Volumes consumed of 1.6 mM saccharin solution or tap water were determined by the mass remaining in separate bottles originally containing 200 g of solution (densities: 0.998 ± 0.003 g/mL tap water; 0.997 ± 0.002 g/mL 1.6 mM saccharin) in a 48 hr period. Preference was determined the difference in consumption between the two liquids, with greater than 50% difference indicating saccharin preference. Male groups sizes for saccharin preference testing were: PSEN2 control = 11 kindled, 11 sham; PSEN2-N141I = 9 kindled, 9 sham; APP control = 6 kindled, 5 sham; APP/PS1 = 6 kindled, 6 sham. Female group sizes for saccharin preference testing were: PSEN2 control = 10 kindled, 8 sham; PSEN2-N141I = 10 kindled, 8 sham; APP control = 7 kindled, 7 sham; APP/PS1 = 5 kindled, 3 sham.

### Barnes Maze Test

The Barnes maze is a land-based spatial learning and memory test for rodents. We used a 7-day testing protocol that consisted of habituation (day 0), acquisition to evaluate learning (days 1-4), and a probe trial to evaluate memory retention (day 7). The habituation day was designed for each mouse to familiarize itself to the maze platform and escape chamber. Each acquisition day consisted of 3 trials/mouse, with >30 minute between each trials, allowing each mouse to learn the location of the escape chamber over 4 days and 12 trials. During the probe trial, the escape chamber was removed from the maze, and each mouse was tested on its ability to recall the escape location for the maze. The latency for the mouse to encounter the escape hole (latency to encounter) and the number of incorrect holes explored (errors) prior to finding the escape chamber were recorded by a blinded investigator during the acquisition and probe trial phases as measures of spatial learning and memory. Male group sizes for Barnes maze testing were: PSEN2 control = 26 kindled, 18 sham; PSEN2-N141I = 15 kindled, 12 sham; APP control = 10 kindled, 11 sham; APP/PS1 = 12 kindled, 9 sham. Female group sizes for Barnes maze testing were: PSEN2 control = 18 kindled, 15 sham; PSEN2-N141I = 16 kindled, 14 sham; APP control = 18 kindled, 17 sham; APP/PS1 = 12 kindled, 10 sham.

### Home Cage Behavioral Phenotyper Test

Single-housed mice were monitored at the end of the behavioral testing period (24 hours after the Barnes maze) in an environmental chamber using a Noldus Phenotyper system equipped with Ethovision to assess circadian activities and exploratory behaviors that may be influenced by the AD-associated genotypes or chronic seizure history^61^. The experiments took place over 67 continuously monitored hours wherein the location, movement, and speed of each single-housed mouse was tracked. Each cage was divided into zones (center, perimeter, food, water, hidden) with the duration in each zone tracked and reported. Continuous behavioral activity measurements were then binned into 1-hour sessions and activity tracked across all zones, with average activity of each bin taken for the light periods (Light 1, Light 2, and Light 3) in comparison with the dark periods (Dark 1, Dark 2, and Dark 3). There were 8 separate Phenotyper cohorts (4 male, 4 female) to achieve a minimum of n=6 fully kindled mice (and sham) with and without each AD genotype. Male group sizes for Phenotyper behavioral analysis were: PSEN2 control = 8 kindled, 8 sham; PSEN2-N141I = 7 kindled, 7 sham; APP control = 8 kindled, 8 sham; APP/PS1 = 8 kindled, 8 sham. Female group sizes for Phenotyper analysis were: PSEN2 control = 7 kindled, 8 sham; PSEN2-N141I = 7 kindled, 8 sham; APP control = 8 kindled, 8 sham; APP/PS1 = 9 kindled, 6 sham.

### Brain Tissue Collection and Processing

Mice were euthanized approximately 1 h after the last seizure threshold stimulation session, and brains were collected and hemi-sected longitudinally. The left-brain hemisphere was flash frozen in 2-methylbutane overlaying dry ice and stored in −80°C but not included in this study; the right brain hemisphere was fixed for 24 h in 4% paraformaldehyde at 4°C, then stored in 30% sucrose with 0.05% sodium azide for cryoprotection and preservation at 4°C for this study. The sucrose-submerged right hemisphere was then flash frozen, before cryosectioning (Leica CM1850) into 30-um thick slices from the dorsal hippocampus (AP from Bregma: −1.58 to −2.10 mm, which includes somatosensory cortex). A total of 20 rostral and 20 caudal hippocampal sections were collected sequentially from each mouse, with 2 sequential sections per brain region/mouse mounted onto SuperFrost slides (Fisher). Slides were stored at −80°C until immunofluorescent processing ^36^.

### Immunofluorescence

Slides from the right brain sections in the seizure threshold cohort were washed with 1X PBS (3 x 5 min) before undergoing antigen retrieval with citrate buffer (#ab93678; Abcam) heated at 95 °C for 10 min, then resting at room temperature for 20 min. GAD65+67-labeled slides antigen retrieval for 20 min at 95 °C, then 40 min at room temperature. The slides were then washed with dH_2_O (3 x 2 min) before being permeabilized with 0.3% Triton X-100 in 1X PBS for 30 min, then incubated with 10% goat serum with 0.03% Triton-X 100 in 1X PBS blocking solution for 2 hr at RT. GAD65/67and PV antibodies were applied to the slides in a 1% goat serum, and 0.03% Triton-X in 1X PBS solution. For EAAT2, GFAP, and 6E10, a different blocking reagent (10% goat serum with 5% Triton-X 100 in 1X PBS) was applied for 2 hr at RT and the antibodies were applied with 5% Goat Serum in 1X PBS. Slides for GAD65/67underwent primary antibody incubation at 4 °C overnight, whereas slides for EAAT2, GFAP, and 6E10 were incubated for 2 hr at RT. The next day, the GAD65/67or EAAT2 slides were washed with 1X PBS for 3 x 5 min and incubated with either anti-rabbit secondary antibody or anti-mouse secondary antibody. For GAD65+67, the secondary antibody was applied in 1% goat serum, 0.03% Triton-X in 1X PBS whereas EAAT2 secondary incubation was conducted in 5% goat serum in 1X PBS. A final wash was done with 1X PBS (3 x 5 min) before the slides were coverslipped with Prolong Gold Antifade with DAPI (#P36934; Thermo Fisher).

Photomicrographs for EAAT2 and GAD65/67 were then captured from subregions of the dorsal hippocampus (CA1, CA3, and DG) and from the somatosensory cortex overlaying CA1 with a fluorescent microscope (Leica DM-6) with a 20x objective (80x final magnification) at minimum 24 hr after coverslip application. Acquisition settings were held constant throughout. GAD65/67 and EAAT2 expression was reported as percent total field area with fluorescent signal for the molecular marker of interest and quantified with ImageJ (Fiji). 6E10 labeling for Aβ was visually confirmed by a blinded investigator in a binary manner (yes/no).

### Statistical Analysis

Seizure thresholds in the minimal clonic and 6 Hz tests were calculated as median convulsant current (CC_50_) and 95% confidence intervals by the probit regression model of Finney^54^, such that statistical difference was defined by non-overlapping confidence intervals (Figure 1). Immunofluorescence field area with EAAT2 or GAD65/67 signal was assessed by t-test between APP control and APP/PS1 mice.

Corneal kindling metrics were assessed by t-test or Mantel Cox Log-Rank tests compared between control and AD mice within each sex. Barnes maze performance and Phenotyper activity were assessed by 3-factor mixed-effects model (time x kindling x genotype), with p<0.05 considered significant (GraphPad Prism v11.0 or later). Differences in saccharin consumption were assessed by 2-way ANOVA Sidak’s multiple comparisons test with the effects of genotype and the effects of kindled status examined.

## Results

### Age-related increases in seizure susceptibility are disrupted in APP/PS1, but not PSEN2-N141I, mice

Seizure threshold in rodents normally increases with advanced age ^62^ and there are known sex-dependent seizure threshold differences^45^. We have earlier reported the age-related changes in minimal clonic and 6 Hz seizure threshold of PSEN2 knockout mice and reported significant divergence from matched controls by 8-months-old ^56^, thus, we aimed to establish whether this phenomenon was differentially apparent in AD models that do (APP/PS1) and do not (PSEN2-N141I) develop Aβ plaques. Minimal clonic seizure threshold was modestly increased with advanced age in both male and female PSEN2-N141I and APP/PS1 mice (Figure 1A-D), but there were no significant and sustained genotype-related differences in seizure threshold. However, there were marked genotype-related differences in seizure threshold in the 6 Hz test in APP/PS1, but not PSEN2-N141I, mice (Figure 1E-H). Notably, only APP/PS1 males (Figure 1G) and females (Figure 1H) showed significant and sustained differences from their sex-matched controls in this test. Notably, the age-related increase in 6 Hz seizure threshold was blunted by 8 months-old, around the time that Aβ plaque deposition becomes widespread in APP/PS1 mice ^63^. This loss of normal age-related 6 Hz seizure threshold increase in APP/PS1 mice clearly demonstrates disruptions in limbic connectivity in this mouse model.

### Cortical and limbic EAAT2 expression is increased in aged APP/PS1 mice

Given that APP/PS1 were significantly more susceptible to an evoked 6 Hz seizure, a mouse model of evoked focal seizures driven by the limbic system^52^, we sought to characterize whether there were notable differences in excitatory (EAAT2) or inhibitory (GAD65/67) marker expression in somatosensory cortex or hippocampal regions of 12-month-old APP/PS1 mice following their last evoked 6 Hz seizure. The human parietal lobe typically shows significant degeneration, loss of grey/white matter, and metabolic decline in AD ^64^, thus we assessed whether there were any changes in EAAT2 or GAD65/67 expression in the mouse somatosensory cortex (Figure 2A and 2E), which closely corresponds to human parietal lobe. There was significantly more EAAT2 expression in aged APP/PS1 mice in somatosensory cortex (t=2.548, p=0.019) and DG (t=2.145, p=0.043) versus matched control mice (Figure 2A and 2D). However, there was no significant difference in GAD65/67 expression in any assessed brain region (Figure 2E-H), indicating that excitatory/inhibitory marker expression was not simply generally increased in these regions in this AD model. Further, we confirmed that aged APP/PS1 mice in our seizure threshold cohort showed the canonical Aβ plaque accumulation in both somatosensory cortex and hippocampus (Figure 2I and 2J), with plaques typically surrounded by reactive astrocytes (arrows in 2I and 2J). Thus, APP/PS1 mice were more hyperexcitable in late life (Figure 1) and this was in part driven by greater EAAT2 expression in cortical and hippocampal structures where Aβ accumulation was also evident. We next assessed how increased neuronal hyperexcitability and chronic seizure history could influence cognitive and functional outcomes in late life.

### Aged AD mouse genotypes are differentially susceptible to corneal kindling

We have previously demonstrated across several different ages and AD-related genotypes that corneal kindling occurs more rapidly in AD-associated models ^24; 42; 56; 65^, and that chronic seizures can influence cognitive function in aged PSEN2 knockout models^36^. However, we had yet to define the interaction between Aβ overexpression and a history of chronic seizures on cognitive and functional outcomes in late life, at ages that more closely approximate human EOAD onset. Consistent with our earlier reports, 12-month-old+ male and female PSEN2-N141I mice did not generally show marked differences in kindling acquisition versus matched controls (Figure 3). However, 12-month-old+ male and female APP/PS1 mice were consistently more prone to corneal kindling than their matched controls, mirroring our earlier studies of pharmacological responses in similarly aged mouse lines^24^. Further, the severity of the kindled seizure was worsened in APP/PS1 male mice (Supplemental Figure 1). Aged fully kindled APP/PS1 and PSEN2-N141I mice then underwent behavioral testing to ascertain the extent to which chronic seizures interact with AD genotypes to influence later life cognitive and behavioral performance.

### Chronic seizures worsen late life cognitive function exclusively in female APP/PS1 mice

Limited clinical evidence suggests that chronic seizures in AD cases may worsen functional decline, including worsened language processing and memory^3^ and survival^66^. Thus, we assessed whether repeated focal seizures could synergistically or additively worsen learning and memory performance in aged male and female PSEN2-N141I and APP/PS1 mice using the land-based Barnes maze task (Figure 4). Measuring the latency to escape into the goal chamber, PSEN2-N141I (Figure 4-B) and APP/PS1 (Figure 4C-D) mice consistently exhibited a time-related improvement, regardless of biological sex. However, there was a time x kindling interaction for male PSEN2-N141I mice (F(4,168)=3.89, p=0.005; Figure 4A), with post-hoc tests showing that performance on the probe trial significantly differed between kindled and sham-kindled mice, regardless of genotype (p<0.05). Male APP/PS1 mice showed a significant time x genotype interaction (F(4, 52)=3.31, p=0.017; Figure 4C), reflecting that APP/PS1 male mice performed consistently poorer than their matched controls but that seizures did not further compound this effect in APP/PS1 mice. However, female APP/PS1 mice were more substantially negatively affected by chronic seizures, showing a significant genotype x kindling interaction on the number of errors to locate the hidden escape chambers (F(1,31)=4.223, p=0.048; Figure 4H). There was a non-significant trend for the probe trial performance to be worse in the kindled APP/PS1 females (p=0.06; Figure 4H). These findings cumulatively suggest that while seizures generally male mouse performance regardless of AD genotype, the cognitive function of female APP/PS1 mice was significantly worsened by seizures on this land-based learning and memory task.

**Figure 4.**
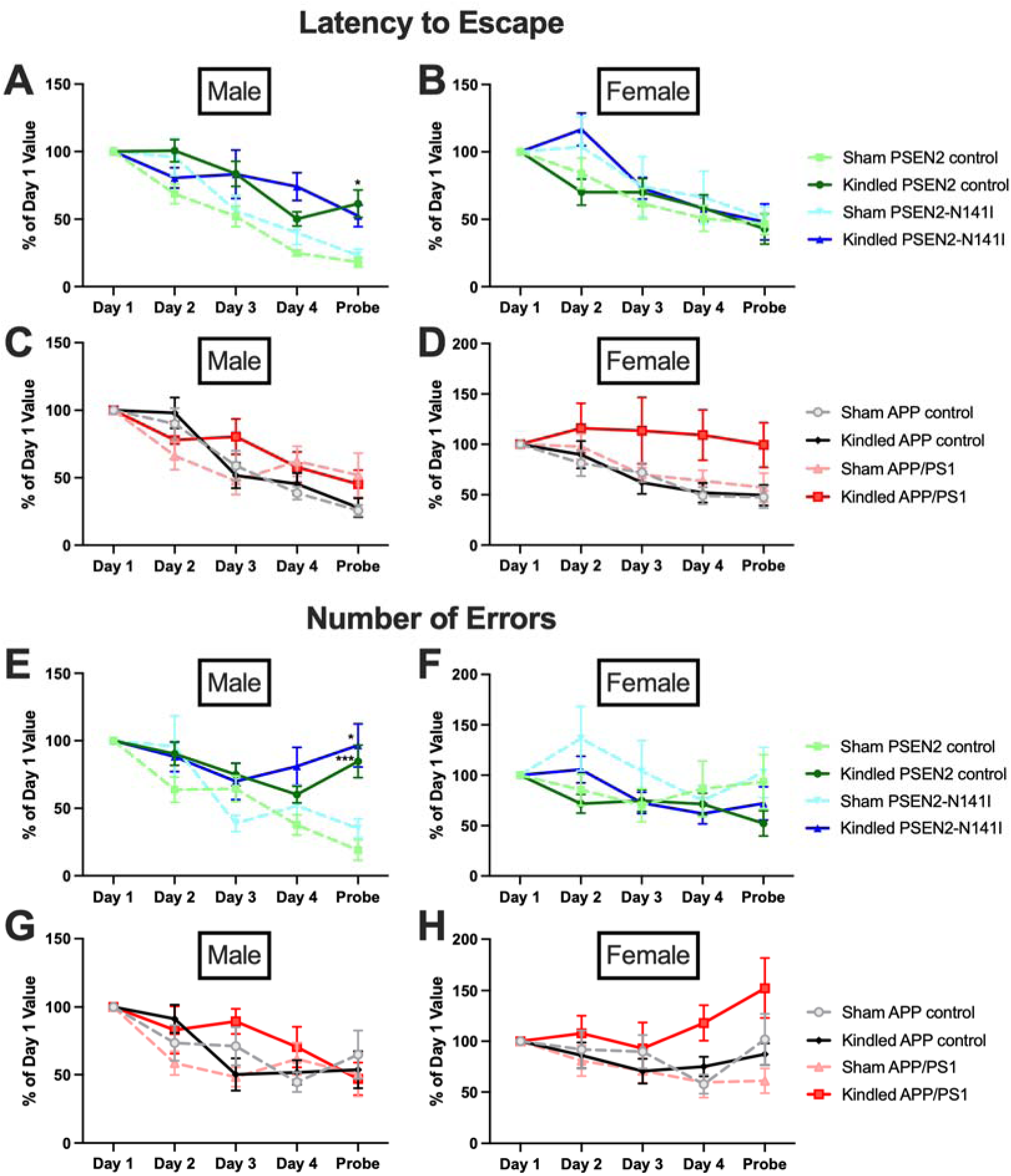
Performance of aged male and female PSEN2-N141I and APP/PS1 mice and their matched genotype control mice with a history of chronic kindled seizures was compared to the performance of animals that did not have a history of chronic seizures using the land-based Barnes maze task of spatial learning and memory. A-D) The latency to escape from the open maze platform was recorded across all sexes and genotypes. A) PSEN2-N141I and genotype matched control male mice demonstrated a significant time x kindling interaction effect, with the final probe trial showing significant differences between kindled and sham kindled mice (time x kindling interaction - F (4, 168) = 3.886, p=0.0048, with post-hoc probe trial p<0.05). B) Female PSEN2-N141I and controls only showed a main effect of time on the Barnes maze task, but neither genotype nor kindling affected performance. C) Male APP/PS1 mice generally performed worse on the Barnes maze task than genotype-matched controls over the testing period, but there was no further worsening by chronic seizures. D) Female APP/PS1 mice generally performed worse on the Barnes maze task (main effect of genotype - F (1, 32) = 4.760, p=0.037), regardless of kindling history. E-H) The number of erroneously explored escape holes was also recorded for all experimental conditions across the testing period. E) Male PSEN2-N141I mice showed a significant time x kindling interaction on Barnes maze performance (F (4, 168) = 7.125, p<0.0001), with the PSEN2-N141I sham mice exhibiting significant differences from genotype-matched kindled mice on the probe test (*p<0.05). Further, sham-kindled PSEN2 control mice significantly differed from kindled PSEN2-control mice (*** p=0.0003). F) Female PSEN2-N141I and PSEN2 control mice generally performed well on the Barnes maze task with regard to errors made, showing only a main effect of time on the task for all experimental groups (F (4, 124) = 3.595, p=0.008). G) Male APP/PS1 and matched control mice showed a similar main effect of time on the Barnes maze task (F (4, 80) = 11.33, p<0.0001), but no genotype or kindling interactions on performance. H) However, female APP/PS1 mice showed a significant kindling x genotype interaction (F (1, 31) = 4.223, p=0.048), suggesting that seizures only affected the cognitive function of these mice in this task. Across all tests, data was compared by 3-factor mixed-effects model (time x kindling x genotype), with p<0.05 considered significant. * indicates p<0.05.

### AD-associated genotypes and chronic seizure history do not worsen depressive-like behaviors

Anhedonia is a feature of clinical depression that can be modeled in the preclinical setting using the saccharin preference test. Anhedonia, but not depression, may also be a significant predictor of conversion to AD in patients with mild cognitive impairment ^67^. Rodent models of epilepsy also demonstrate notable reductions in saccharin preference ^68^. Further, PSEN conditional double KO mice demonstrate more significant age-related neuropsychiatric deficits, e.g. anxiety, irritability, depression, apathy, aggressivity, and anhedonia when compared with other AD-like mouse models ^69^, suggesting that presenilins may more meaningfully contribute to the comorbid neuropsychiatric comorbidities of AD. Therefore, to define whether chronic seizures worsen AD-associated neuropsychiatric comorbidities in an Aβ-dependent manner, we quantified the saccharin preference of kindled and sham-kindled PSEN2-N141 and APP/PS1 mice versus their matched controls (Figure 5). The combination of documented seizure history and AD-related genotypes did not differentially impact saccharin preference in either sex, suggesting no additive or synergistic interaction of chronic seizures on this neuropsychiatric comorbidity of AD.

**Figure 5.**
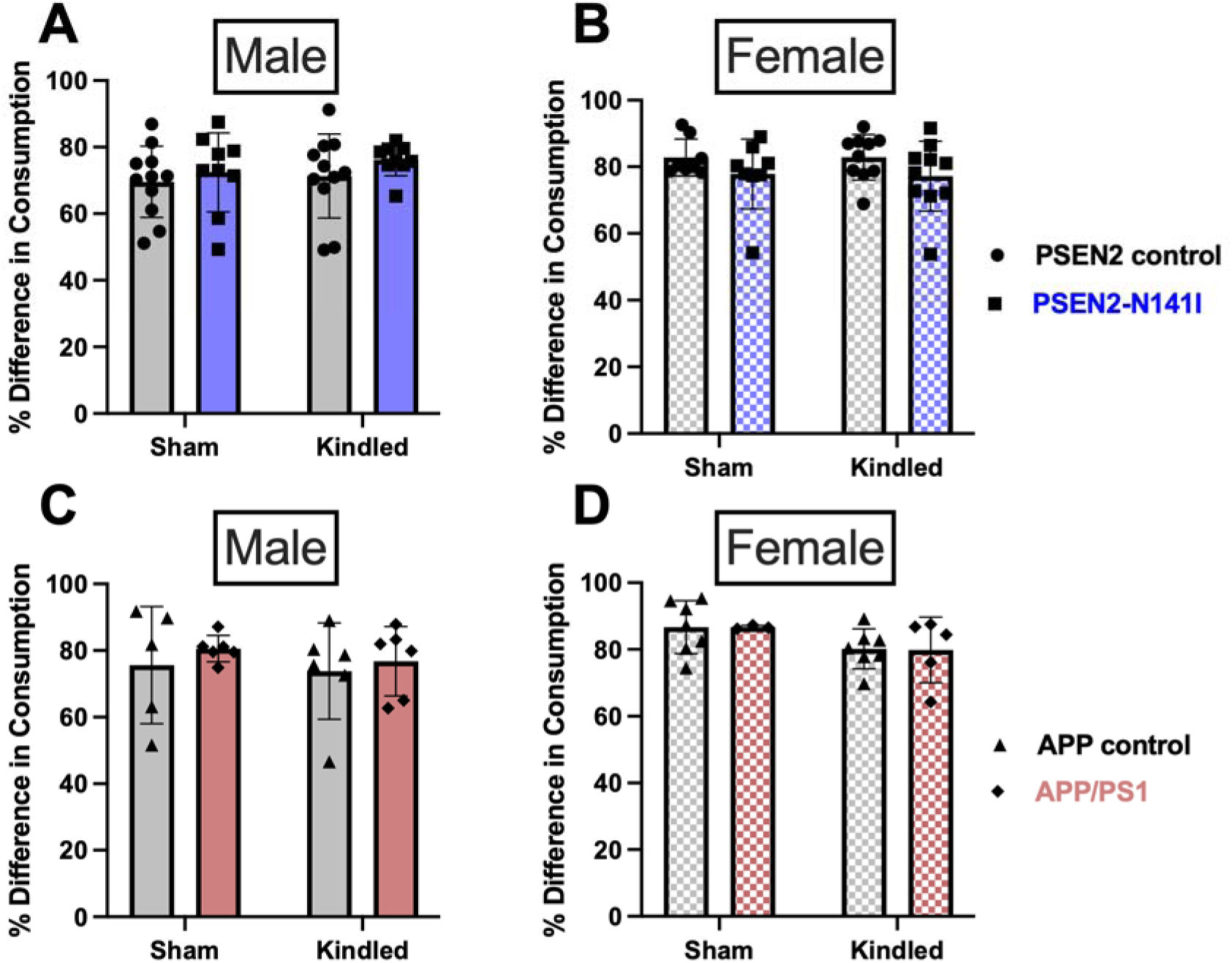
Aged male and female AD-associated mice with and without a history of chronic kindled seizures do not show anhedonia-like behavior as assessed by the saccharin preference test (% Difference in Consumption = saccharin solution consumed (g) / total liquid consumed (g)). Neither the PSEN2-N141I genotype nor a history of chronic kindled seizures significantly influenced the level of saccharin consumption in either A) male or B) female aged mice. There was also no change in saccharin consumption levels in C) male or D) female aged APP/PS1 mice compared to genotype-matched controls. Differences in saccharin consumption were assessed by 2-way ANOVA Sidak’s multiple comparisons test with the effects of genotype and the effects of kindled status examined.

### Circadian behavior patterns are disrupted by AD-related genotypes in a sex-specific manner

Patients with AD often report sleep disturbances, possibly due to brain hyperactivity thus we assessed whether chronic seizure affects mouse behavior in a genotype- or sex-specific manner over a 67-hour continuous home cage monitoring session (Figure 6 and Figure 7). Male APP/PS1 mice showed consistent time x kindling x genotype interactions across multiple assessed longitudinal domains, including total exploratory behavior (F(66, 922)=1.87, p<0.0001; Figure 6B), frequency in the center zone (F(66, 923)=1.336, p=0.042; Figure 6D), frequency in the perimeter zone (F(66,924)=1.70, p<0.0006; Figure 6F), total duration in the perimeter zone (F(66,924)=1.671, p<0.0009; Figure 6H), and total duration in the hidden zone (i.e. a hut-style nest, F(66,924)=1.71, p=0.0005; Figure 6J). Conversely, male PSEN2-N141I mice were less likely to be affected by chronic seizures, only demonstrating a time x kindling interaction on total activity during the monitoring period (F(66,858)=1.63, p=0.0024; Figure 6A) and a genotype x kindling interaction on frequency in the perimeter zone (F(1,13)=8.28; p=0.013; Figure 6E). Thus, the home cage circadian activities of male PSEN2-N141I and APP/PS1 were differentially impacted by chronic seizures.

**Figure 6.**
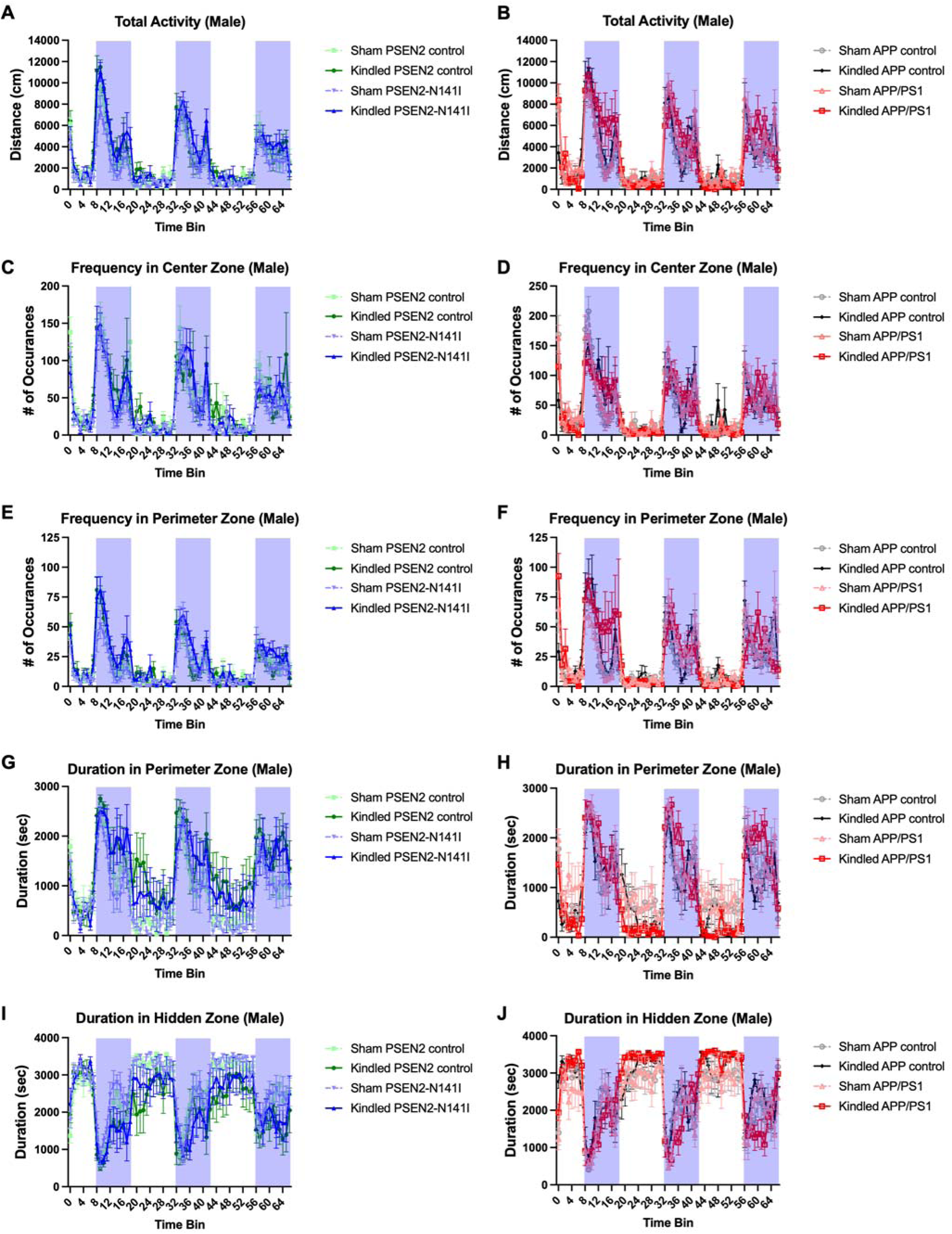
Chronic evoked seizures dramatically alter circadian behaviors in male APP/PS1, but not PSEN2-N141I, mouse AD models. All analyses were measured by 3-factor mixed-effects model and Tukey’s post-hoc test (n=4/group). A) Total activity in the entire chamber during each 1-hour bin for a 67-hour recording session for PSEN2-N141I and PSEN2 control male mice. There was a significant time x kindling group interaction (F (66,789) = 1.617, p=0.0019). B) Total activity in the entire chamber during each 1-hour bin for a 67-hour recording session for APP/PS1 and APP/PS1 control male mice. There was a significant time x kindling group interaction (F (66,922) = 1.87, p<0.0001). Frequency in the center zone during each 1-hour bin for a 67-hour recording session for male C) PSEN2-N141I and D) APP/PS1 mice was similarly assessed. There was a significant main effect of time for PSEN2-N141I male mice (C), but a significant time x kindling x genotype interaction for APP/PS1 mice (D). Frequency in the perimeter zone during each 1-hour bin for a 67-hour recording session for male E) PSEN2-N141I and F) APP/PS1 mice was similarly assessed. There was a significant genotype x kindling effect in PSEN2-N141I male mice (E), but a significant time x kindling x genotype interaction for APP/PS1 mice (F). Duration in the perimeter zone during each 1-hour bin for a 67-hour recording session for male G) PSEN2-N141I and H) APP/PS1 mice was similarly assessed. There was a significant main kindling effect in PSEN2-N141I male mice (G), but a significant time x kindling x genotype interaction for APP/PS1 mice (H). Duration in the hidden zone (i.e. a plastic “hut”) during each 1-hour bin for a 67-hour recording session for male G) PSEN2-N141I and H) APP/PS1 mice was similarly assessed. There was a significant main time effect in PSEN2-N141I male mice (G), but a significant time x kindling x genotype interaction for APP/PS1 mice (H). Altogether, these findings demonstrate discrete genotype specific effects on circadian behavioral patterns in AD-related mouse models over the course of 67 hours of continuous recording.

Female PSEN2-N141I and APP/PS1 mice showed a similar pattern of differential, genotype-specific seizure-related impacts on circadian behaviors (Figure 7). Specifically, female PSEN2-N141I mice exhibited time x kindling interaction (F(66,789)=2.01, p<0.0001) and a time x genotype interaction (F(66,789)=1.617, p=0.002) on their total home cage activity (Figure 7A), whereas female APP/PS1 mice demonstrated a time x kindling x genotype interaction (F(66, 788)=1.00, p=0.0004; Figure 7B). Kindled APP/PS1 female mice were consistently less active in the dark cycle relative to all other experimental conditions (Figure 7B). Frequency in the center zone of the home cage was consistently impacted with a time x kindling interaction in both PSEN2-N141I (F(66,782)=1.682, p=0.0009; Figure 7C) and APP/PS1 (F(66,789)=2.17, p<0.0001; Figure 7D).

**Figure 7.**
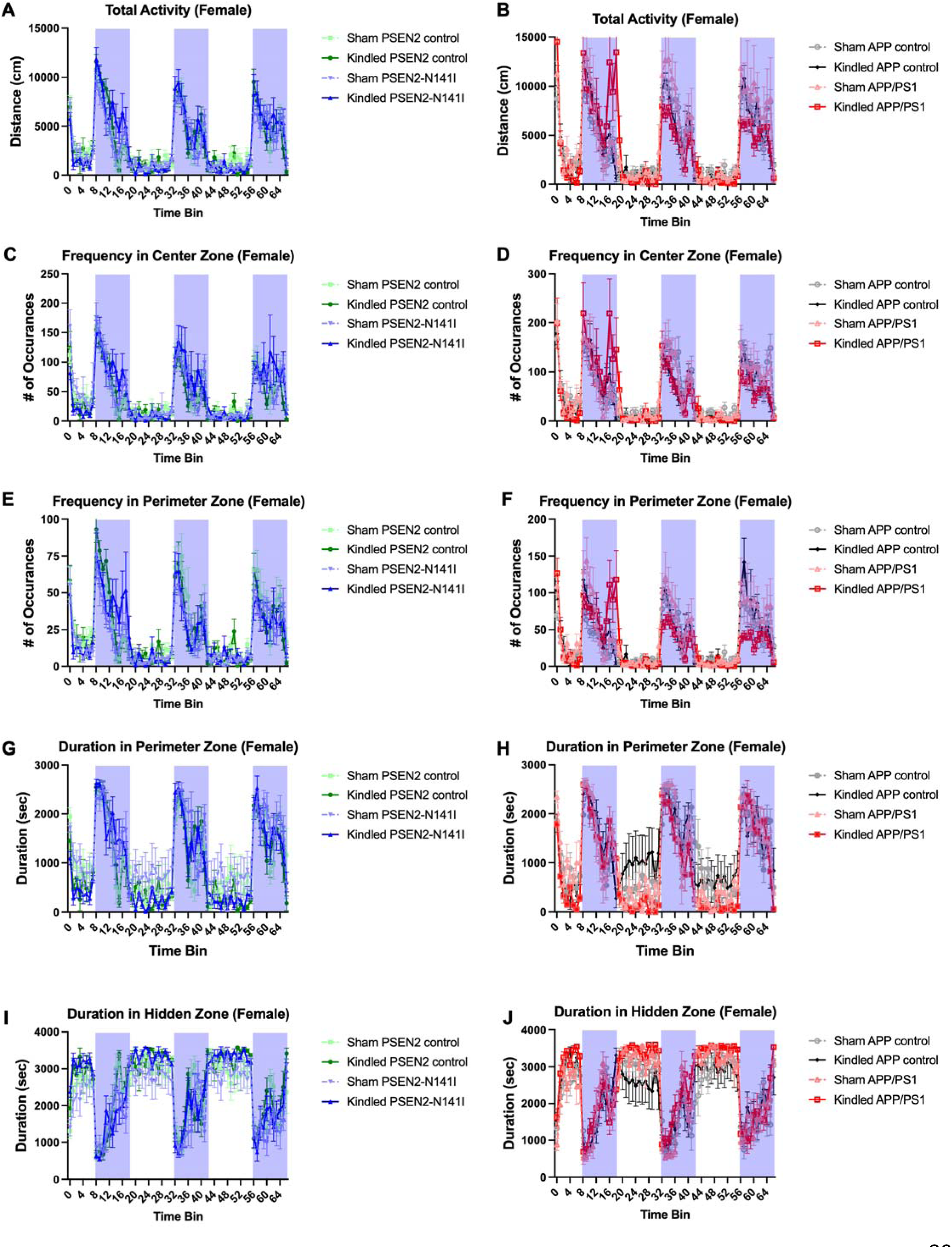
Chronic evoked seizures dramatically alter circadian behaviors in both female PSEN2-N141I and APP/PS1 mouse AD models. All analyses were measured by 3-factor mixed-effects model and Tukey’s post-hoc test (n=4/group). A) Total activity in the entire chamber during each 1-hour bin for a 67-hour recording session for PSEN2-N141I and PSEN2 control female mice. There was a significant time x kindling group interaction and a significant time x genotype interaction. B) Total activity in the entire chamber during each 1-hour bin for a 67-hour recording session for APP/PS1 and APP/PS1 control female mice. There was a significant time x kindling group interaction (F (66,788) = 1.00, p=0.0004). Frequency in the center zone during each 1-hour bin for a 67-hour recording session for female C) PSEN2-N141I and D) APP/PS1 mice was similarly assessed. There was a significant time x kindling effect of time for both female PSEN2-N141I mice (C) and APP/PS1 mice (D). Frequency in the perimeter zone during each 1-hour bin for a 67-hour recording session for female E) PSEN2-N141I and F) APP/PS1 mice was similarly assessed. There was a significant time x kindling effect and a significant time x genotype effect in PSEN2-N141I female mice (E), but a significant time x kindling x genotype interaction for female APP/PS1 mice (F). Duration in the perimeter zone during each 1-hour bin for a 67-hour recording session for female G) PSEN2-N141I and H) APP/PS1 mice was similarly assessed. There was a significant main time x kindling effect in PSEN2-N141I female mice (G), but a significant time x genotype interaction for APP/PS1 female mice (H). Duration in the hidden zone (i.e. a plastic “hut”) during each 1-hour bin for a 67-hour recording session for female G) PSEN2-N141I and H) APP/PS1 mice was similarly assessed. There was a significant main time x kindling interaction effect in PSEN2-N141I female mice (G), but a significant time x kindling x genotype interaction for female APP/PS1 mice (H). Altogether, these findings demonstrate discrete genotype specific effects on circadian behavioral patterns in AD-related mouse models over the course of 67 hours of continuous recording.

However, there was both a time x genotype interaction (F(66,790)=1.656, p=0.0012) and a time x kindling interaction (F(66,790)=1.344, p=0.040) on the frequency in the perimeter zone in PSEN2-N141I female mice (Figure 7E), whereas female APP/PS1 mice showed a time x kindling x genotype interaction (F(66,789)=2.40, p<0.0001; Figure 7F). Duration in the perimeter zone was impacted with a time x kindling interaction in female PSEN2-N141I mice (F(66,790)=1.713, p=0.0006; Figure 7G), but in a time x genotype related interaction manner in female APP/PS1 mice (F(66,786)=1.784, p<0.0001; Figure 7H). Finally, the duration in the hidden zone (i.e. a “hut”) showed a kindling x time interaction in PSEN2-N141I female mice (F(66,790)=1.803, p=0.0002; Figure 7I) but a time x genotype interaction in APP/PS1 female mice (F(66,786)=1.69, p=0.0008; Figure 7J). Notably, female APP/PS1 mice showed a non-significant trend for a time x genotype x kindling interaction on the duration in the hidden zone (F(66,786)=1.25, p=0.096; Figure 7J). Cumulatively, these data spanning 67 hours of home cage monitoring in male and female AD-related mice demonstrate that chronic seizures differentially influence circadian behaviors of aged mice with AD-related genetic risk factors in a sex- and genotype-specific manner. Our present study with APP/PS1 versus PSEN2-N141I mice clearly demonstrates that APP overexpression may differentially impact epileptogenesis and AD burden in late life, providing compelling mechanistic insight into the urgency to address the focal seizures in AD patients as a novel interventional approach to disease modification.

We also assessed exploratory behavior of PSEN2-N141I in a 10 minute OF chamber exploration chamber task to determine whether chronic seizure history influenced anxiety-like behaviors in male and female mice (Supplemental Figure 2). Kindling generally increased exploratory behavior of male and female PSEN2-N141I mice (Supplemental Figure 2), but there were no significant kindling x genotype interactions on the OF task. To further establish whether longer duration monitoring in the Phenotyper home cage monitoring paradigm could differentially reveal genotype-related effects of seizure history, we also quantified the total center zone exploration in each of the genotypes during the first light cycle tracking period (6 hours of recording; Supplemental Figure 3). Chronic kindled seizures evoked in male APP/PS1 and matched control mice consistently reduced center zone preference in the Phenotyper chambers (Supplemental Figure 3B; F(1,28)=19.67, p<0.0001), whereas PSEN2-N141I male mice were not significantly affected (Supplemental Figure 3A). Conversely, female PSEN2-N141I and their matched controls with chronic kindled seizures showed generally reduced chamber center zone exploration over this longer duration paradigm versus sham-kindled mice (PSEN2-N141I – F(1,25)=17.04; p=0.0004; Supplemental Figure 3C). This trend was also evident in APP/PS1 and matched control female mice (APP/PS1 – F(1,25)=15.03, p=0.0006; Supplemental Figure 3D). Thus, longer duration home cage monitoring revealed consistent sex-related differential impacts of chronic seizures, but no significant genotype x seizure interactions on total exploration in the light cycle.

## Discussion

This study provides critical direct evidence of the functional and behavioral impacts of untreated repeated secondarily generalized focal seizures in the setting of both advanced age and discrete AD-associated genetic risk factors. Most notably, our study reveals significant heterogeneity in the consequences of evoked seizures across multiple behavioral aspects of AD, including spatial working memory and circadian patterns of behavior in both male and female mice. This differential impact of chronic evoked seizures in aged animals provides invaluable insight into the discrete clinical domains that may be functionally impacted by untreated focal seizures in AD. Our study provides compelling justification that the early treatment of focal seizures may be a reasonable disease modification strategy to beneficially slow AD progression.

Prior studies of seizures in AD mouse models have largely focused on outcomes associated with SRS and electrographic activity passively recorded in relatively young, single-housed male mice that overexpress Aβ ^12; 13^, leaving unanswered significant questions related to the direct contributions of seizures on AD processes in late life. We have previously established the corneal kindling model in a variety of AD-related mouse lines to assess the direct contributions of seizures to long-term behavioral and functional outcomes ^36; 42^ and pharmacological antiseizure drug efficacy^24; 56^, thus we aimed to extend these prior studies to experimentally assess the functional impact of investigator-initiated chronic focal seizures on neuropsychiatric comorbidities (saccharin preference test), spatial learning and memory (Barnes maze), circadian activity (Phenotypers), and pathological changes (immunofluorescence) in aged PSEN2-N141I and APP/PS1 mice using age ranges that closely approximate clinical EOAD. Notably, our present study utilized the heterozygous PSEN2-N141I mouse to model the neuronal and non-neuronal biology of human AD patients with this gene variant ^20; 70^. This study is strengthened because it juxtaposed this Aβ-independent mouse model against the impacts of seizures in an amyloidogenic mouse model (APP/PS1), thereby allowing us to directly interrogate the additive or synergistic contribution that APP overexpression, and thus Aβ accumulation, has on the development of neuropsychiatric sequelae and neuropathology in the context of chronic seizures versus other AD-related pathological features such as neuroinflammation. This head-to-head comparison of the impact of chronic seizures in PSEN2-N141I versus APP/PS1 mice allowed us to discretely interrogate the role that Aβ-dependent versus Aβ-independent mechanisms may play in combination with chronic focal seizures to impact the severity of neuropsychiatric deficits, cognitive dysfunction, and neuropathology in late life.

Age-related changes in seizure susceptibility are clinically well-documented and can be reproduced in rodent epilepsy models. While epilepsy tends to be most common in the very young and the very old ^41; 71; 72^, little insight is available to understand how seizure susceptibility changes across the lifespan in individuals harboring AD-related risk genes. Thus, we first sought to establish the changes, if any, in seizure susceptibility with advanced age in PSEN2-N141I and APP/PS1 mice. While minimal clonic seizure threshold was generally not different between AD models and their matched controls, mirroring our prior findings in PSEN2 knockout mice ^56^, meaningful genotype-specific differences emerged in the 6 Hz test. The 6 Hz seizure test is a minimally invasive electrically evoked acute seizure model that can establish the integrity and efficiency of neuronal circuits in a rodent with genetic variants^45; 56^. These seizures mimic the focal seizures of human temporal lobe epilepsy and tend to engage very different brain regions from other evoked seizure paradigms, such as the maximal electroshock and subcutaneous pentylenetetrazol test ^52^. In neurologically normal or seizure-naïve animals, the necessary mechanisms for forebrain-gated seizure onset function separately, but with genetic manipulation, such as the PSEN2 variant or APP overexpression presently employed, this neurocircuitry becomes interconnected and the neuronal network efficiency is improved, leading to reduced focal seizure threshold. Notably, this present study demonstrates that the normal age-related increases in 6 Hz seizure threshold in both male and female APP/PS1 is significantly blunted relative to age-matched WT littermates, likely coinciding with the age of Aβ plaque accumulation beginning at around 4.5-6 months-old ^63^. Further, there were no significant changes in 6 Hz seizure threshold in PSEN2-N141I mice versus their WT counterparts, further suggesting a role for Aβ accumulation on reducing seizure threshold. These findings suggest that while AD genotypes do alter clinical seizure risk and patients with the PSEN2-N141I variant are more prone to clinical seizures ^20^, the integrity and connectivity of the neurocircuitry underlying focal seizure generation is likely not adversely affected with loss of normal PSEN2 function, whereas the presence of Aβ may adversely influence this neurocircuitry.

In this study, we demonstrate that EAAT2 expression is significantly altered in the somatosensory cortex and hippocampal DG of APP/PS1 mice aged >12 months-old, two brain regions known to be dysfunctional in human epilepsy and AD. The AD brain exhibits neuroimaging and neurophysiology changes that may contribute to clinical seizures and neuronal hyperexcitability ^73^; our present observations of enhanced EAAT2 expression in the somatosensory cortex, a brain region closely tied to the human parietal cortex, may contribute to this aberrant neurocircuitry. Moreover, the observation that GAD65/67expression was not altered indicates that this was not a global change in its expression in APP/PS1 mice. These findings altogether suggest that APP/PS1 mice are more seizure prone in late life, thus compelling us to further explore how chronic seizures in this genotype impacted cognitive function and neuropsychiatric comorbidities.

Focal seizures and epileptiform activity are consistently documented in AD cases ^1; 74^ and animal models ^12; 13; 75–77^. However, recording spontaneous epileptiform activity or overt focal SRS in mouse models is labor-intensive and costly, requiring a tremendous resource investment to obtain sufficient statistical power to discern the *in vivo* effects of infrequent spontaneous seizures. In this regard, moderate- to high-throughput models of evoked focal seizures are critical to more efficiently and directly assess the biological impact of unchecked neuronal hyperexcitability on AD burden and progression. The gold-standard in epilepsy research is the recording of spontaneous epileptiform discharges and SRS; however, EEG is impractical for sufficiently powered behavioral studies with costly aged rodents. Furthermore, social isolation stress can dramatically lower seizure threshold in rodents, leaving already seizure-prone animals at heightened risk for SRS, which may further complicate study interpretation. In this regard, the corneal kindling model allows for robust investigation of the direct effects of chronic seizures on disease outcomes in an investigator-controlled manner that retains social housing throughout the seizure monitoring period. Further, just as in humans with seizures and dementia ^78^, we find that chronic kindled seizures evoke premature mortality in young and aged APP/PS1 mice^24; 42^, further validating this approach for AD studies. Therefore, we adapted a moderate-throughput corneal kindled seizure test to AD-associated mouse models to differentiate the biological response to evoked seizures for this present study. Notably, the mouse corneal kindled mouse test is one of the most clinically predictive preclinical screening models for focal-onset seizures in people with epilepsy ^79; 80^ and is highly sensitive to levetiracetam ^81^, an antiseizure medicine (ASM) presently in clinical trial for AD, further underscoring this seizure model’s relevance to AD modification interventional studies.

Evoked 60 Hz kindled seizures in young APP/PS1, but not PSEN2-N141I, mice increase Aβ production^42^ and APP/PS1 mice with 6 Hz corneal kindled seizures also show increased Aβ plaque levels relative to WT controls^65^. Our present studies altogether provide additional evidence that excess neuronal hyperexcitability may incite the onset of AD pathology (i.e., increase Aβ levels), suggesting that Aβ plaque deposition is a functional consequence of this hyperexcitability-initiated pathological cascade, rather than a cause ^42^. While Aβ-lowering therapies have been approved for clinical AD, they also pose safety risks ^82–84^ and offer modest therapeutic benefit to slow AD progression. In contrast, treatment with ASMs are effective and well-tolerated in older adults with epilepsy, and emerging clinical evidence suggests that AD outcomes may be improved with ASM administration in the prodromal period ^2; 85^. Thus, our present study provides compelling direct evidence that seizures in AD may altogether worsen critical behavioral domains, including sex- and genotype-specific detrimental effects to cognitive function and circadian behaviors, such that ASM treatment to reduce focal seizure activity may be a tractable disease-modifying strategy for AD. Numerous ASMs are approved for people with epilepsy, and these medications offer a promising untapped avenue that should be further mechanistically pursued to expedite drug repurposing for disease modification in AD. Further, given that seizures accelerate cognitive and functional decline in AD ^2^, and that current Aβ-lowering treatments offer limited efficacy at the expense of safety, addressing and managing the seizures symptoms in AD could significantly reduce the estimated $190B annual U.S. healthcare burden associated with this neurodegenerative disease.

---

**Supplemental Figure 1.**
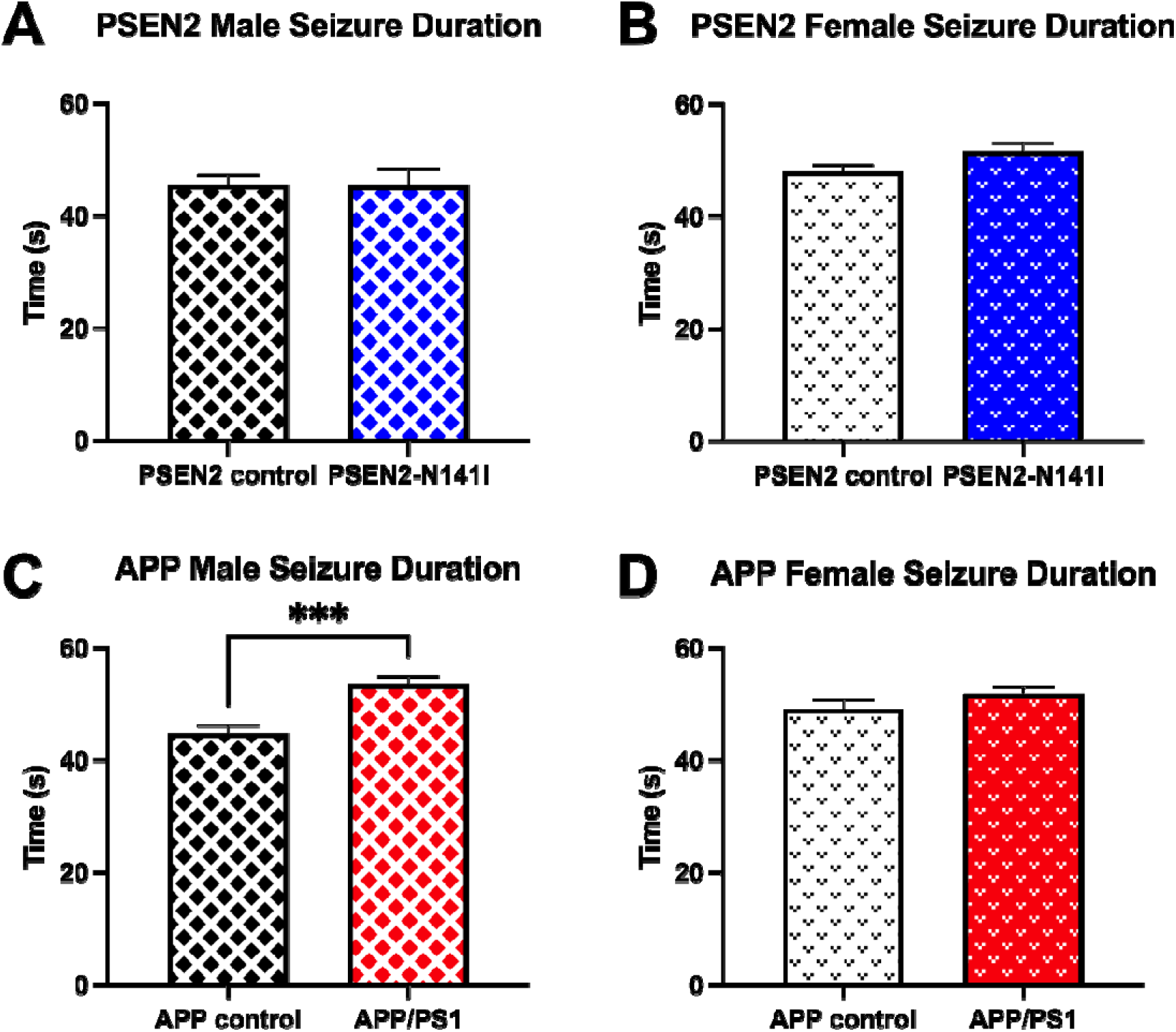
Genotype significantly affects kindled seizure duration in fully-kindled male mice. A, B) The PSEN2-N141I genotype did not significantly influence the post-kindling seizure length in aged male or female (p=0.07) mice. C) Aged APP/PS1 male mice had a significantly increased average seizure length compared to APP controls (p=0.0002). D) There was no effect of genotype on seizure duration in the aged APP/PS1 females when compared to APP controls. Differences in seizure duration means were assessed by unpaired t test. *** = p<0.001.

**Supplemental Figure 2.**
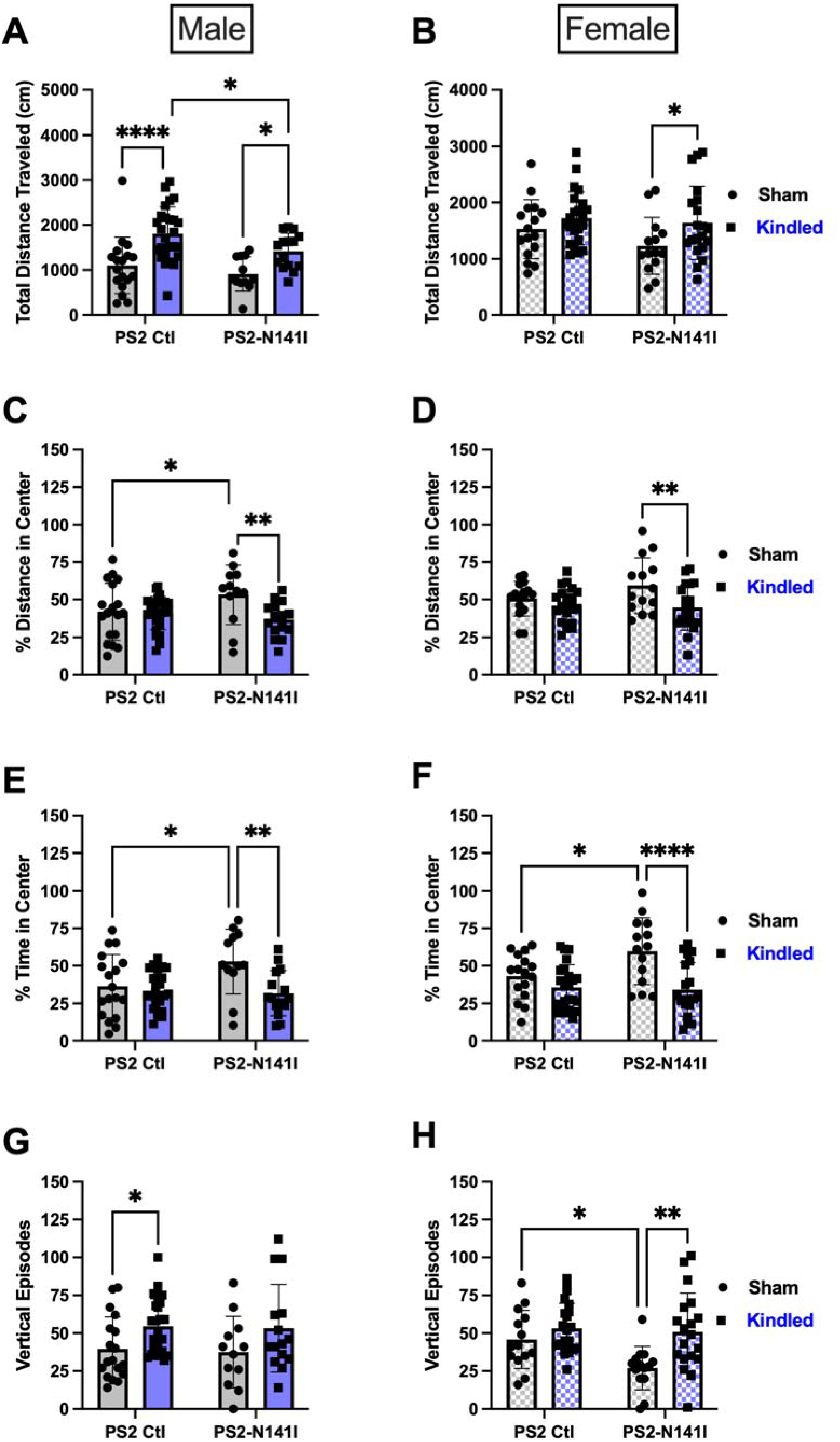
The PSEN2-N141I genotype did not impact overall motor activity or anxiety-like behavior in aged mice in an open field paradigm. A) Kindling significantly increased the distance traveled in the open field in male control animals compared to shams (p=0.0003), a slightly diminished effect was seen in the PSEN2-N141I kindled males compared to shams (p=0.07). B) There was no effect of kindling status or genotype on distance traveled by female PSEN2-N141I mice or controls. C-D) Vertical rearing episodes, a sign of exploration, were increased in female kindled PSEN2-N141I mice compared to sham-kindled PSEN2-N141I mice (p=0.005). (E-F) Sham PSEN2-N141I of each sex traveled a higher percentage of their total distance in the center of the open field than kindled male (p=0.03) and female (p=0.02) PSEN2-N141I mice. (G-H) Sham PSEN2-N141I mice also spent more time in the center of the open field than kindled PSEN2-N141I mice (male p=0.02; female p=0.0006). Differences in open field activity were assessed by 2-way ANOVA Sidak’s multiple comparisons test with the effects of genotype and the effects of kindled status examined. * = p<0.05, ** = p<0.01, *** = p<0.001.

**Supplemental Figure 3.**
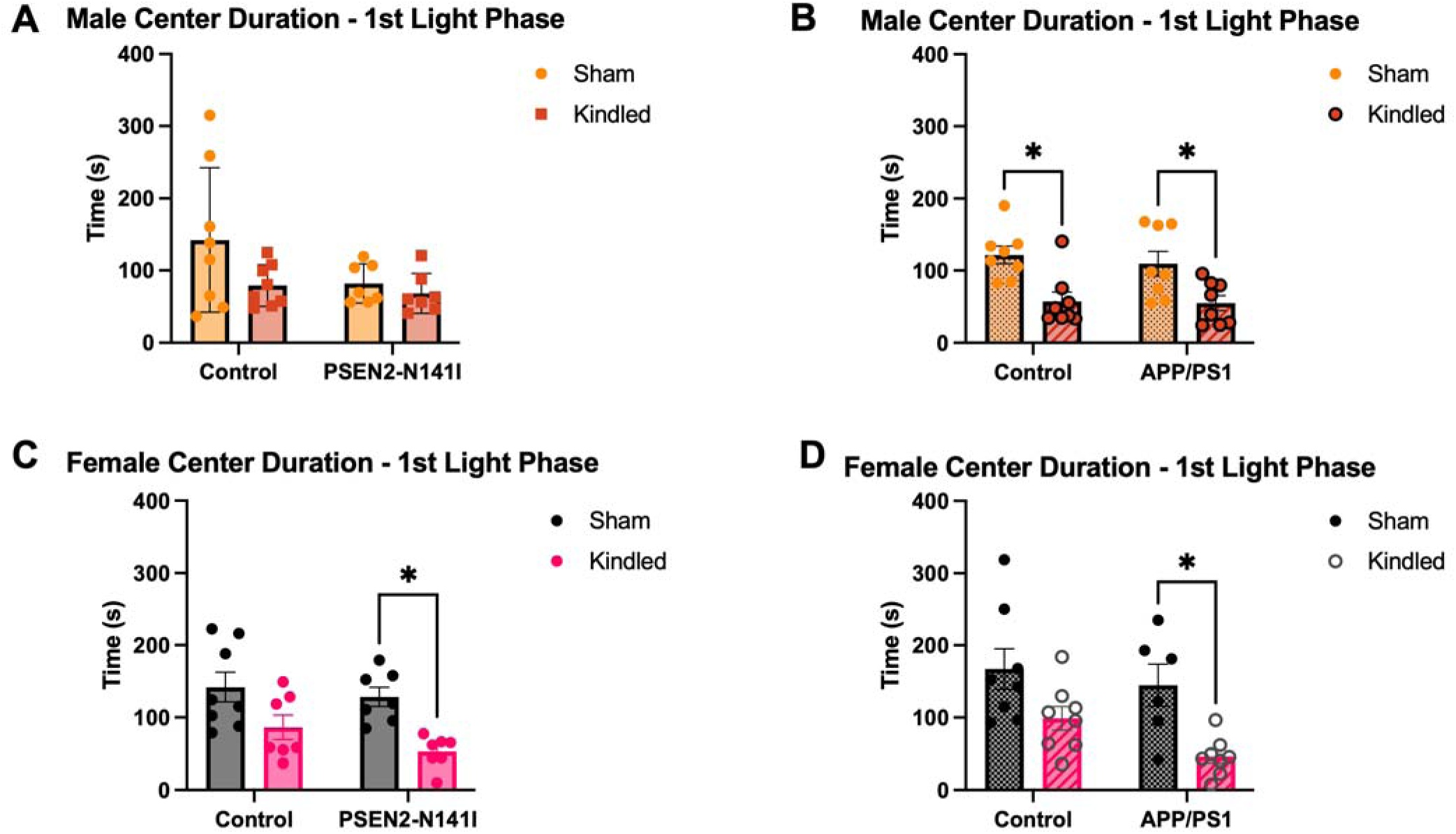
Chronic evoked seizures lead to changes in home cage exploration over a 67-hour continuous monitoring period in a Phenotyper chamber. A) Male PSEN2-N141I and matched controls did not show significant exploration differences with chronic seizure history. B) Male APP/PS1 and APP/PS1 control mice showed a main effect of kindling on center duration explored in the home cage (F(1,28)=19.67, *p<0.0001). C) Female PSEN2-N141I and matched controls explored the Phenotyper chamber less with chronic kindled seizure history (F(1,25)=17.04, p=0.0004). D) Female APP/PS1 and matched controls explored the Phenotyper chamber less with chronic kindled seizure history (F(1,25)=15.03, p=0.0006).

## Acknowledgements

Research reported in this publication was supported by an American Epilepsy Society Young Investigator Award, the University of Washington School of Pharmacy, and the NIA of the National Institutes of Health under grant number R01AG067788. A total of 50% of the project was financed with federal funds. A total of 50% percent of the project was financed with non-governmental funds. The authors acknowledge the technical support of Leanne Lehmann, Ali Lo, and Rami Koutoubi.

