## Supplemental Figure 1 for "Chronic evoked seizures differentially worsen cognitive function and long-term neuropsychiatric comorbidities in Alzheimer’s disease-related genotypes"

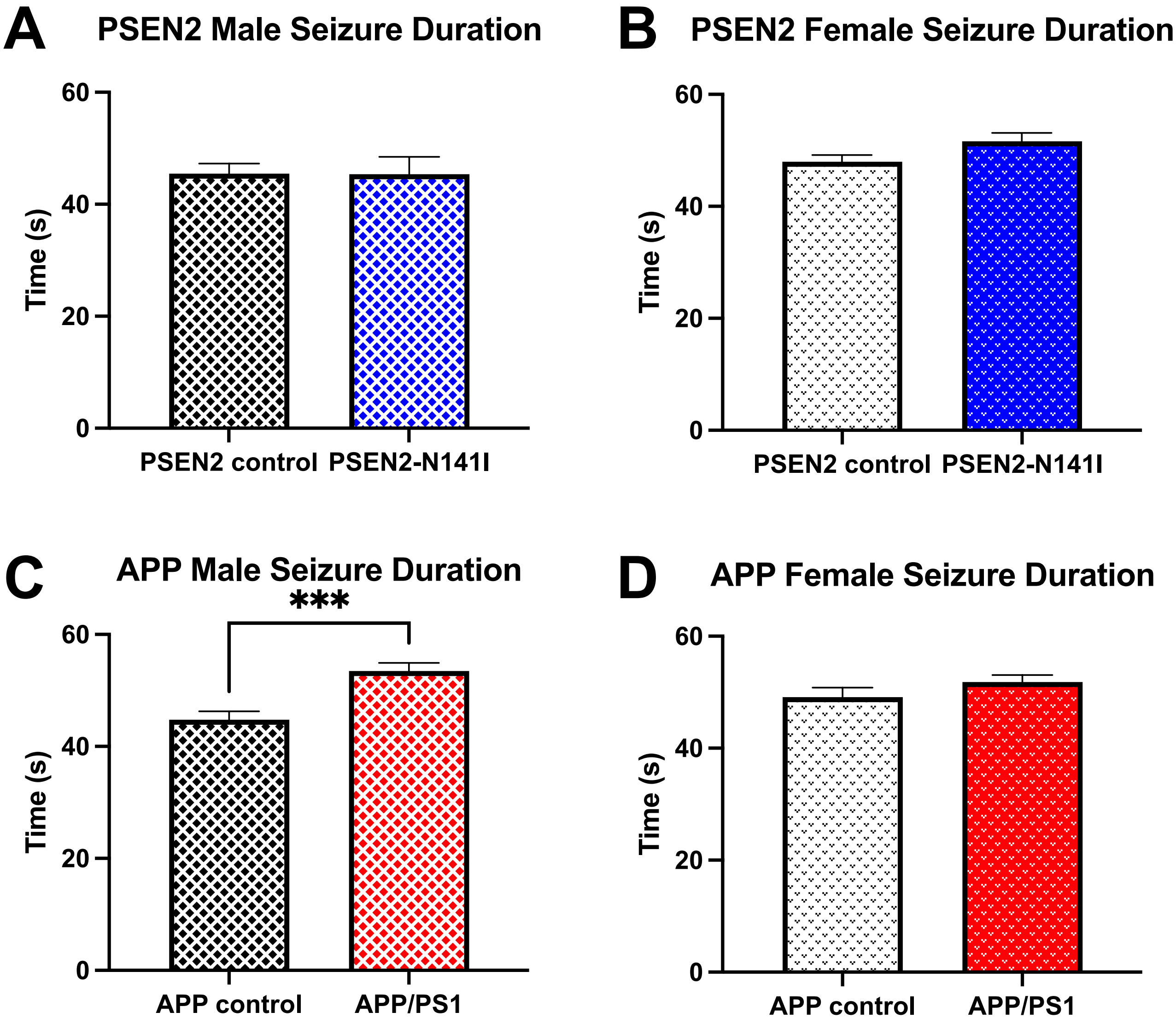
A
PSEN2 Male Seizure Duration
Time (s)
0
20
40
60
PSEN2 control
PSEN2-N141I
B
PSEN2 Female Seizure Duration
Time (s)
0
20
40
60
PSEN2 control
PSEN2-N141I
C
APP Male Seizure Duration
***
Time (s)
0
20
40
60
APP control
APP/PS1
D
APP Female Seizure Duration
Time (s)
0
20
40
60
APP control
APP/PS1
