## Supplementary figures and images for "Chronic evoked seizures differentially worsen cognitive function and long-term neuropsychiatric comorbidities in Alzheimer’s disease-related genotypes"

### Supplemental Figure 2

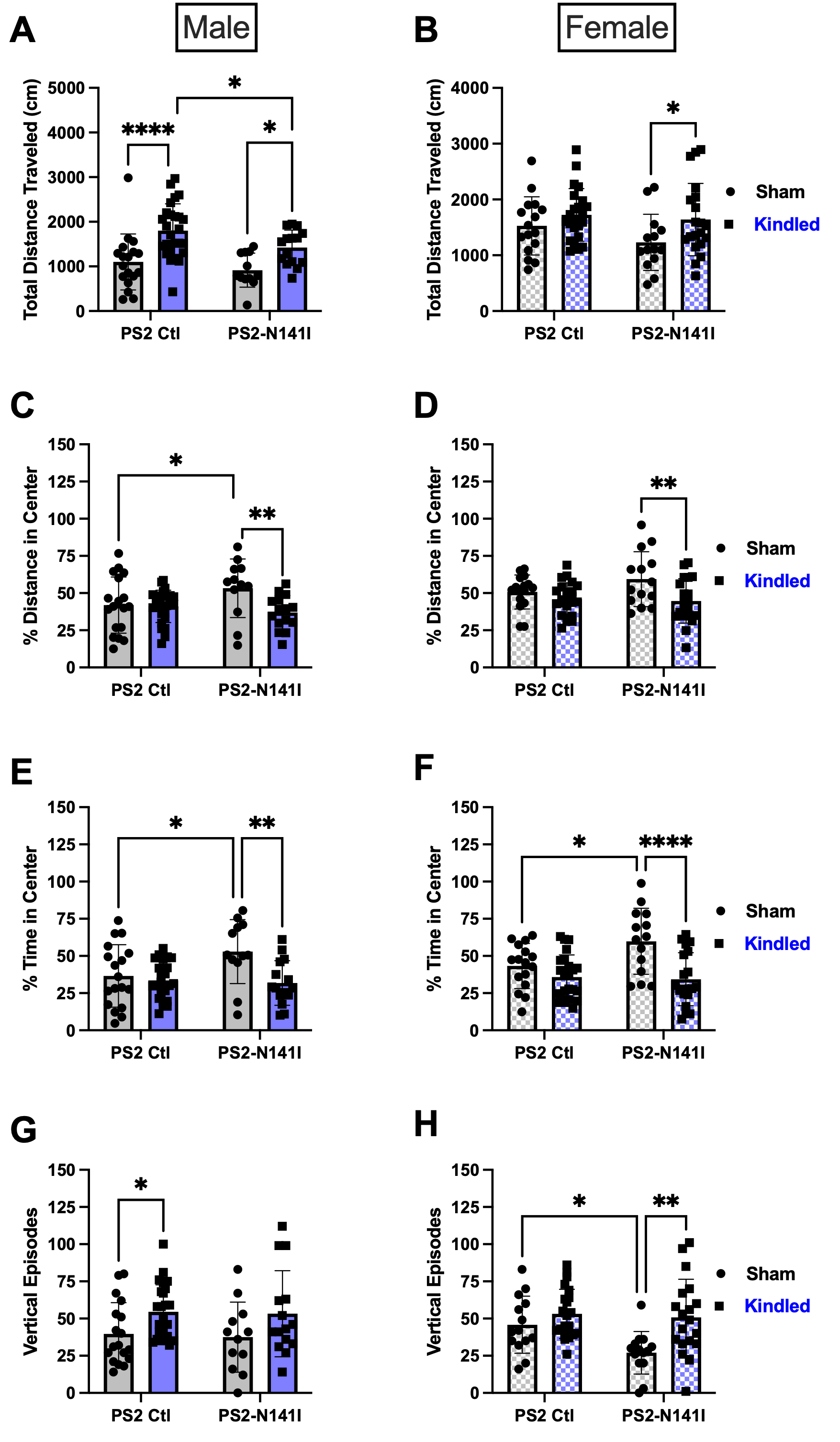

### Supplemental Figure 3

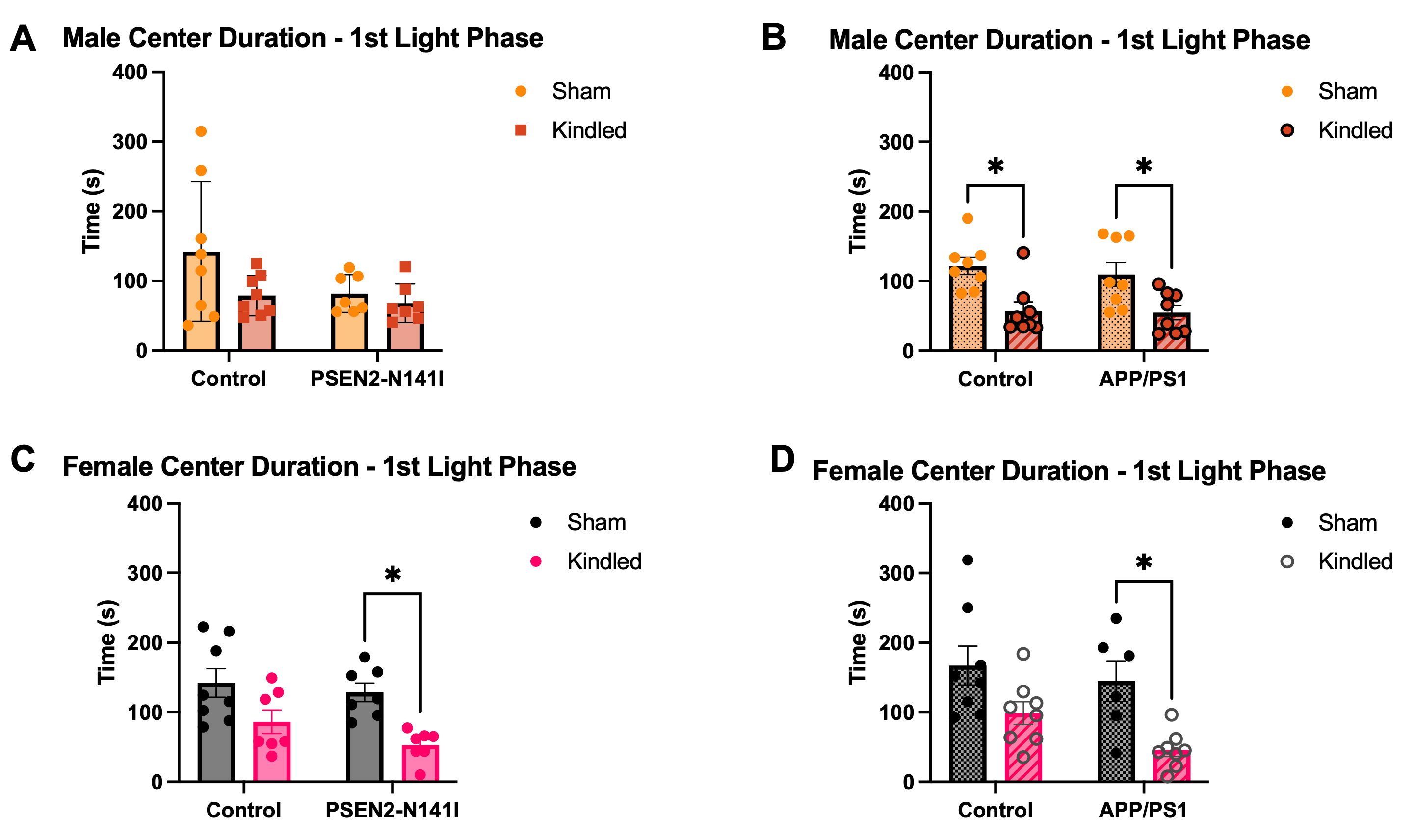
